# Differential gene expression elicited by ZIKV infection in trophoblasts from congenital Zika syndrome discordant twins

**DOI:** 10.1101/867465

**Authors:** Murilo Sena Amaral, Ernesto Goulart, Luiz Carlos Caires-Júnior, David Abraham Morales-Vicente, Alessandra Soares-Schanoski, Roselane Paiva Gomes, Giovanna Gonçalves de Oliveira Olberg, Renato Mancini Astray, Jorge E. Kalil, Mayana Zatz, Sergio Verjovski-Almeida

## Abstract

Zika virus (ZIKV) causes congenital Zika syndrome (CZS), which is characterized by fetal demise, microcephaly and other abnormalities. ZIKV in the pregnant woman circulation must cross the placental barrier that includes fetal endothelial cells and trophoblasts, in order to reach the fetus. CZS occurs in ∼1-40% of cases of pregnant women infected by ZIKV, suggesting that mothers’ infection by ZIKV during pregnancy is not deterministic for CZS phenotype in the fetus. Therefore, other susceptibility factors might be involved, including the host genetic background. We have previously shown that in three pairs of dizygotic twins discordant for CZS, neural progenitor cells (NPCs) from the CZS-affected twins presented differential *in vitro* ZIKV susceptibility compared with NPCs from the non-affected. Here, we analyzed human-induced-pluripotent-stem-cell-derived (hiPSC-derived) trophoblasts from these twins and compared by RNA-Seq the trophoblasts from CZS-affected and non-affected twins. Following *in vitro* exposure to a Brazilian ZIKV strain (ZIKV^BR^), trophoblasts from CZS-affected twins were significantly more susceptible to ZIKV^BR^ infection when compared with trophoblasts from the non-affected. Transcriptome profiling revealed no differences in gene expression levels of ZIKV candidate attachment factors, IFN receptors and IFN in the trophoblasts, either before or after ZIKV^BR^ infection. Most importantly, ZIKV^BR^ infection caused, only in the trophoblasts from CZS-affected twins, the downregulation of genes related to extracellular matrix organization and to leukocyte activation, which are important for trophoblast adhesion and immune response activation. In addition, only trophoblasts from non-affected twins secreted significantly increased amounts of chemokines RANTES/CCL5 and IP10 after infection with ZIKV^BR^. Overall, our results showed that trophoblasts from non-affected twins have the ability to more efficiently activate genes that are known to play important roles in cell adhesion and in triggering the immune response to ZIKV infection in the placenta, and this may contribute to predict protection from ZIKV dissemination into fetuses’ tissues.

**Author summary:** The Zika virus (ZIKV) infection in adults is usually characterized by mild flu-like symptoms, with most cases remaining asymptomatic. However, in the last years, widespread ZIKV infection was shown for the first time to be associated with congenital Zika syndrome (CZS) and death of neonates. CZS is a very debilitating condition that includes microcephaly and mental retardation, leading to a strong social and health impact. This dramatic condition calls for a careful evaluation of the molecular mechanisms involved in ZIKV infection in the maternal-fetal interface. It is estimated that CZS occurs in ∼1-40% of cases of pregnant women infected by ZIKV, which suggests that different susceptibility factors might be involved, including the host genetic background. By analyzing trophoblast cells that recapitulate the placenta from three pairs of dizygotic twins discordant for CZS, we were able to show that trophoblasts from CZS-affected twins were significantly more susceptible to ZIKV infection when compared with trophoblasts from the non-affected twins. We also provide a detailed picture of genes differentially expressed by trophoblasts from the discordant twins after infection with ZIKV. These genes can be further investigated as possible therapeutic targets to avoid viral dissemination into developing fetus’ tissues.

## Introduction

Zika virus (ZIKV) is a flavivirus with sporadic outbreaks reported in several countries, causing an infection usually characterized by mild symptoms, where up to 80% of cases remain asymptomatic [1-3]. However, most likely due to the mutation acquired during the large outbreak recorded in French Polynesia in 2013–2014 [4-6], for the first time, widespread ZIKV infection was shown to be associated with congenital Zika syndrome (CZS) and death of neonates [3,7-13].

CZS is characterized by variable clinical presentations, including fetal demise, microcephaly and other abnormalities (hearing and ocular loss, mental retardation, epilepsy, muscle weakness, learning disabilities and behavioral abnormalities) [14-16]. It is well established now that ZIKV shows tropism for a wide range of host tissues [17-19], especially neuronal cell types, including neural progenitor cells, mature neurons and astrocytes (reviewed by Christian et al. [20]), and replicates in *ex vivo* slices from adult human cortical tissues [21]. Also, Zika virus infection reprograms global transcription in the host cells [22]. ZIKV in the maternal circulation needs to cross the placental barrier that includes fetal endothelial cells and trophoblasts in order to reach the fetus [23,24]. Several placenta related cells have been shown to be infected by ZIKV, including placental macrophages and trophoblasts [25-29].

It has been estimated that CZS occurs in ∼1-40% of cases of pregnant women infected by ZIKV [3,13,30,31]. This suggests that mothers’ infection by ZIKV during pregnancy is not the only factor determining CZS phenotype in the fetus, and other susceptibility factors might be involved. Indeed, neural progenitor cells (NPCs) from different individuals have been shown to respond differently to ZIKV infection [32,33]. In this scenario, discordant twins represent a good case–control sample to test for the genetic contribution determining the fetuses’ discordant outcome of gestational infection with ZIKV, as they are supposed to have been exposed to ZIKV under similar conditions in the uterus during gestation.

We have previously shown that twins discordant for CZS outcome show differential neural progenitor cells (NPCs) *in vitro* viral susceptibility to a Brazilian ZIKV strain (ZIKV^BR^) [32], but it remains to be determined if discordant CZS twins also show differential placental susceptibility to viral infection. Here, we compared the susceptibility and molecular signatures associated with *in vitro* ZIKV^BR^ infection of trophoblasts from the same CZS-affected and non-affected twins, using a well-established trophoblasts model that recapitulates the primitive placenta formed during implantation [34,35]. We show here that hiPSC-derived trophoblasts from CZS-affected twins were significantly more susceptible to *in vitro* ZIKV^BR^ infection when compared with trophoblasts from non-affected twins. In this context, we had previously shown that, before infection, ESC-derived trophoblasts express a wide range of attachment factors for ZIKV entry and lack the components of a robust antiviral response system [35]; in contrast, cells from term placentas, which resist infection, do not express genes encoding attachment factors implicated in ZIKV entry and do express many genes associated with antiviral defense [35]. However, no ZIKV infection assays were performed with these ESC-derived trophoblasts [35]. Here, transcriptome profiling of hiPSC-derived trophoblasts revealed that ZIKV^BR^ infection elicited different responses in hiPSC-derived trophoblasts from CZS-affected and non-affected twins, highlighting that genes involved with extracellular matrix organization as well as with immune response activation in the placental tissue may contribute to modulate ZIKV infection outcome.

## Results

### Infection with ZIKV^BR^ of hiPSC-derived trophoblasts from discordant twins

We obtained blood from three pairs of dizygotic twins discordant for CZS (non-affected: #10608-4, #10763-4, and #10788-4; CZS-affected: #10608-1, #10763-1, and #10788-1) for generation of hiPSC-derived trophoblasts and for phenotypic and gene expression analysis after *in vitro* infection with ZIKV^BR^ (**Fig 1A**). Erythroblasts from the three pairs of DZ twins were reprogrammed towards hiPSCs. All hiPSC lines were previously shown by immunofluorescence staining to express markers of pluripotency (TRA-1-60 and OCT4) and by RT-qPCR to express endogenous pluripotent transcription factors including *NANOG* and *OCT4* [32].

**Fig 1.**
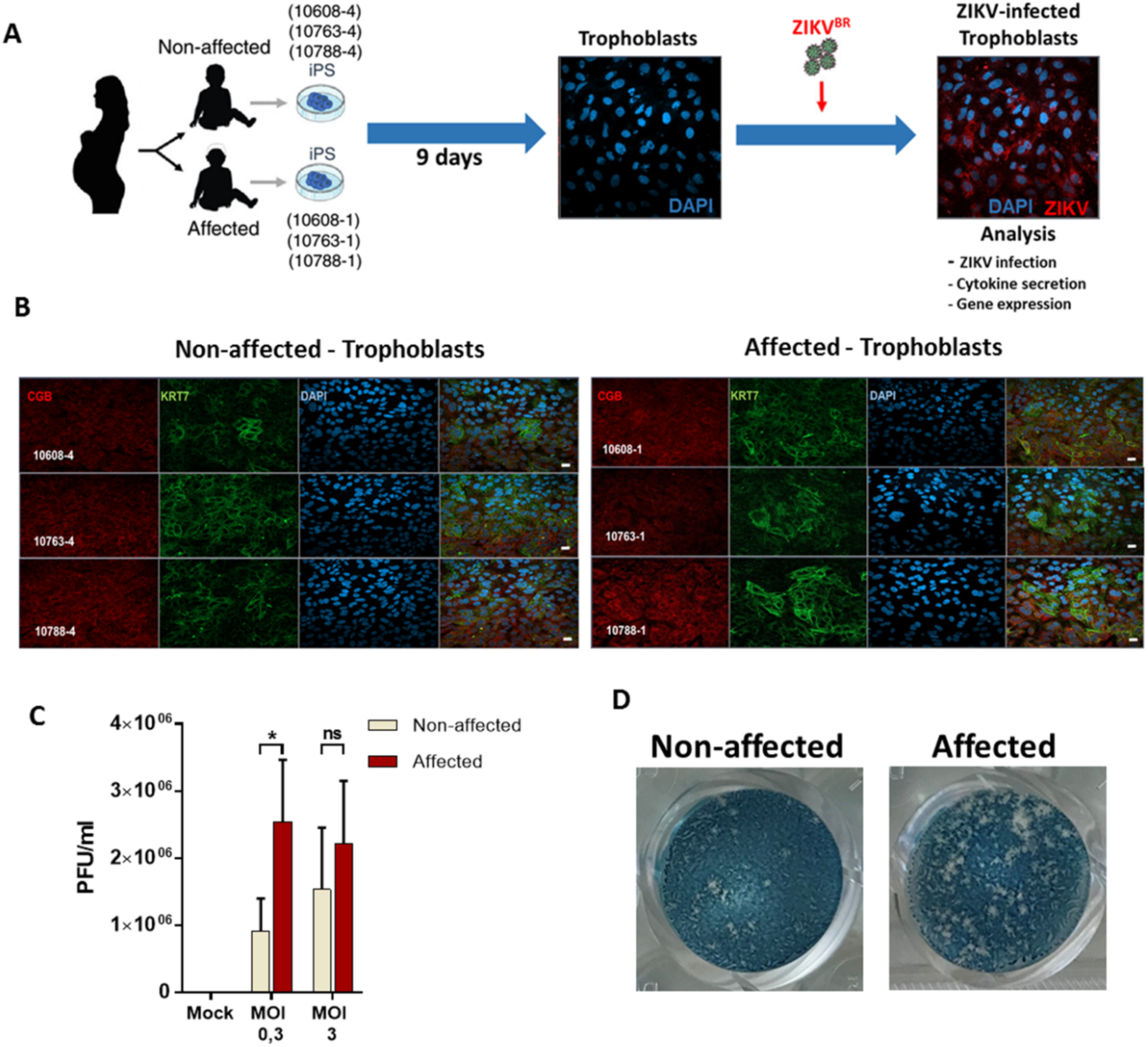
Experimental design and ZIKV^BR^ infection in hiPSC-derived trophoblasts. **(A)**. Schematic: generation of trophoblasts from congenital Zika syndrome affected and non-affected discordant twins’ hiPSCs followed by ZIKV^BR^ infection and analysis. Silhouettes are courtesy of www.vecteezy.com (mother) and Yulia Ryabokon (babies). (**B)**. Immunofluorescence for (β-CG, human chorionic gonadotropin β) and KRT7 (CK7, cytokeratin 7, a pan trophoblast marker) in hiPSC-derived trophoblasts. Scale bar: 20 μm. (**C)**. ZIKV PFU/mL in trophoblasts’ supernatant at MOI = 0.3 and MOI = 3; mean ± SEM of the three twins; *p < 0.05; Student’s t test. (**D)**. Representative plaque forming assay wells with stained VERO cells exposed to ZIKV collected at 96 hpi from the culture supernatants of affected or non-affected #10608 twins’ hiPSC-derived trophoblasts infected at a MOI of 0.3.

Then, the hiPSCs originated from the three pairs of twins were differentiated into primitive trophoblasts using an established protocol [34,35] and further characterized to confirm their differentiation *in vitro*. Under these conditions, we found that all hiPSC-derived trophoblast lines robustly expressed chorionic gonadotropin subunit beta 3 (β-CG/ hCGB3) and keratin 7 (CK7/KRT7) (**Fig 1B**), two of the most commonly used trophoblast markers [36].

Subsequently, we infected hiPSC-derived trophoblasts with ZIKV^BR^ using multiplicity of infection (MOI) of 0.3 and 3, and at 96 hpi (hours post-infection) we investigated the viral titer by measuring the number of plaque-forming units (PFU) in cell culture supernatants (**Fig 1C**). Virus titers were significantly higher (2.8-fold) in the supernatant of CZS-affected twins’ trophoblasts infected with MOI 0.3 (**Fig 1C** and **1D**), indicating that trophoblasts from CZS-affected twins were significantly more susceptible to ZIKV infection or at least more virus productive when compared with trophoblasts from non-affected twins. Infection with the higher MOI (MOI = 3) obliterated this difference due to the high susceptibility of trophoblasts in general to the Zika infection. In order to have a better resolution of the differences between cells derived from CZS-affected and non-affected twins, subsequent studies were performed with MOI of 0.3.

### Potential flavivirus attachment factors and IFN receptor genes were not differentially expressed between trophoblasts from discordant twins

To evaluate the possible differences in molecular signatures associated with ZIKV^BR^ infection in trophoblasts from CZS-affected and non-affected twins, we performed RNA-Seq analysis in hiPSC-derived trophoblasts before and after *in vitro* infection with ZIKV^BR^ for 96 h (MOI = 0.3). With this approach, we could analyze possible differences in gene expression of potential attachment factors for ZIKV and of genes related to antiviral response.

We first confirmed the efficiency of hiPSC differentiation into trophoblasts by looking at the expression levels of a set of over100 genes which have been associated with the trophoblast lineage of mammals [35,36]. When compared with hiPSCs, most of these genes showed significant upregulation in trophoblasts after differentiation (**S1 Table**). We validated by RT-qPCR the differential expression of *NANOG*, a hiPSC marker, and of *HCGA, HCGB* and *KRT7*, three of the genes upregulated in the trophoblasts upon differentiation, as compared with the hiPSCs (**S1 Fig**). Consistent with this and with previous experiments that used the same differentiation protocol [35], proliferation-related genes were downregulated in the trophoblasts relative to the hiPSCs (**S2 Fig**). In addition, four genes encoding transcription factors – *CDX2, ELF5, EOMES* and *ASCL1*, which are generally regarded as markers of trophoblast stem cells [37] – were found to be barely expressed (TPM < 1) in all six hiPSC-derived trophoblast samples from the twins analyzed here. This suggests that all trophoblasts from all the twins differentiated beyond the trophoblast stem-cell stage.

We then looked for differences in the expression levels of potential flavivirus attachment factor genes, of IFN genes and of IFN receptor genes between the trophoblasts from CZS-affected and non-affected twins. These differences could be related to the greater susceptibility to ZIKV^BR^ infection of the trophoblasts from CZS-affected twins.

Many potential flavivirus attachment factors have been proposed as ZIKV candidate receptors, including glycosaminoglycans (CD209/DC-SIGN and HSPG*2*) and TAM receptors (TYRO3, AXL, MERTK) [38-40]. None of these receptor genes was found to be differentially expressed between the trophoblasts from CZS-affected and non-affected twins (**S3 Fig**). *CD209/DC-SIGN* gene was barely expressed in the hiPSC-derived trophoblasts either before or after ZIKV^BR^ infection (**S3 Fig**). *HSPG2* gene had the highest expression levels in hiPSC-derived trophoblasts among the genes encoding potential ZIKV candidate receptors, however no differences in the expression levels of *HSPG2* gene between trophoblasts from CZS-affected and non-affected twins were found either before or after ZIKV^BR^ infection (**S3 Fig**). Recently, AXL has been reported as the receptor involved in ZIKV entry in many cell types, including neural stem cells [41], human umbilical vein endothelial cells (HUVECs) [42], primary human astrocytes [43] and Sertoli cells [44]. *AXL* gene expression showed a marked increase upon differentiation from hiPSC towards trophoblasts (> 5-fold on average), but infected and non-infected hiPSC-derived trophoblasts from non-affected or CZS-affected twins show similar expression levels of *AXL* gene (**S3 Fig**). TAM-family receptors bind phosphatidylserine indirectly, through the soluble intermediates GAS6 (growth arrest-specific 6) and PROS1 [45]. Recently, it was shown that ZIKV infects HUVECs much more efficiently than other flaviviruses because it binds GAS6 more avidly, which in turn facilitates its interaction with AXL [42]. Infected and non-infected hiPSC-derived trophoblasts from non-affected or CZS-affected twins show similar expression levels of *GAS6* and *PROS1* genes (**S3 Fig**). HAVCR1 (TIM1; hepatitis C receptor) was shown to have an important role in the entry of ZIKV into primary cell types from mid- and late-gestation placentas and explants from first-trimester chorionic villi [26]. Interestingly, *HAVCR1* gene was barely expressed in all the hiPSC-derived trophoblasts (**S3 Fig**). Expression levels of all the above receptor genes measured here in non-infected hiPSC-derived trophoblasts from the three pairs of twins confirmed previous data from ESC-derived non-infected trophoblasts [35].

Production of interferons has been reported as a key step in the antiviral immune response to ZIKV [25,46,47]. In agreement with Sheridan et al. [35], primitive trophoblasts expressed low levels of mRNAs from all the IFN genes (**S4 Fig**) when compared with ZIKV potential attachment factors (**S3 Fig**). Upon *in vitro* infection with ZIKV^BR^, a marked induction of mRNA expression of *IFNB1, IFNL1, IFNL2* and *IFNL3* genes but not of *IFNE* was observed (**S4 Fig**). Of note, IFN genes were not differentially expressed between the trophoblasts from CZS-affected and non-affected twins (**S4 Fig**). *IFNA, IFNG, IFNK* and *IFNW1* genes were not detectable in any hiPSC-derived trophoblast cells.

Interestingly, *IFNGR1* and *IFNGR2* (type II IFN receptor genes) were highly upregulated in hiPSC-derived trophoblasts when compared with the hiPSCs that originated them (**S5 Fig**). These two IFN receptor genes were not differentially expressed between trophoblasts from CZS-affected and non-affected twins, and their expression was not changed upon ZIKV^BR^ infection (**S5 Fig**). Compared with the expression of type II IFNR genes, the expression levels of the other interferon receptor genes, namely type I IFNR (*IFNAR1* and *IFNAR2*) and the first (*IFNLR1*) and second (*IL10RB*) subunits of type III IFNR gene (*IFNL*) were lower in both hiPSCs and hiPSC-derived trophoblasts (**S5 Fig**), and again none of them had their expression affected by ZIKV^BR^.

### Induction of interferon-stimulated genes (ISGs) expression and of IFN secretion in hiPSC-derived trophoblasts upon ZIKV^BR^ infection

It is known that interferons participate as mediators of the response to ZIKV infection in the maternal-fetal interface [25,48,49] and that ESC-derived trophoblasts lack components of a robust antiviral response system before infection with ZIKV [35]. However, it is not known if hiPSC-derived primitive trophoblasts are able to induce an antiviral response after infection with ZIKV. To determine this, we analyzed our RNA-Seq data and looked for differentially expressed genes (DEGs) in the hiPSC-derived primitive trophoblasts after *in vitro* infection with ZIKV^BR^.

An important gene expression response was induced in both hiPSC-derived trophoblasts from CZS-affected and non-affected twins following *in vitro* infection with ZIKV^BR^, and a set of 471 DEGs were upregulated in hiPSC-derived trophoblasts after infection with ZIKV^BR^ (FDR < 0.05, Edge R exact test) (**Fig 2A**). These genes (**S2 Table**) include interferon-stimulated genes (ISG) and genes related to cytokine secretion. We found a significant (FDR < 0.05, cumulative hypergeometric distribution) enrichment of different Gene Ontology (GO) terms among these 471 upregulated DEGs detected in the RNA-Seq experiment, being the top categories “response to type I interferon”, “defense response to virus” and “negative regulation of viral process” (**Fig 2B, S3 Table**). Consistent with the GO analysis, Ingenuity Pathway Analysis (IPA) also pointed to an enriched network of interferon-stimulated genes upregulated in hiPSC-derived trophoblasts after ZIKV infection (upregulated ISGs are colored in pink in **Fig 3A**).

**Fig 2.**
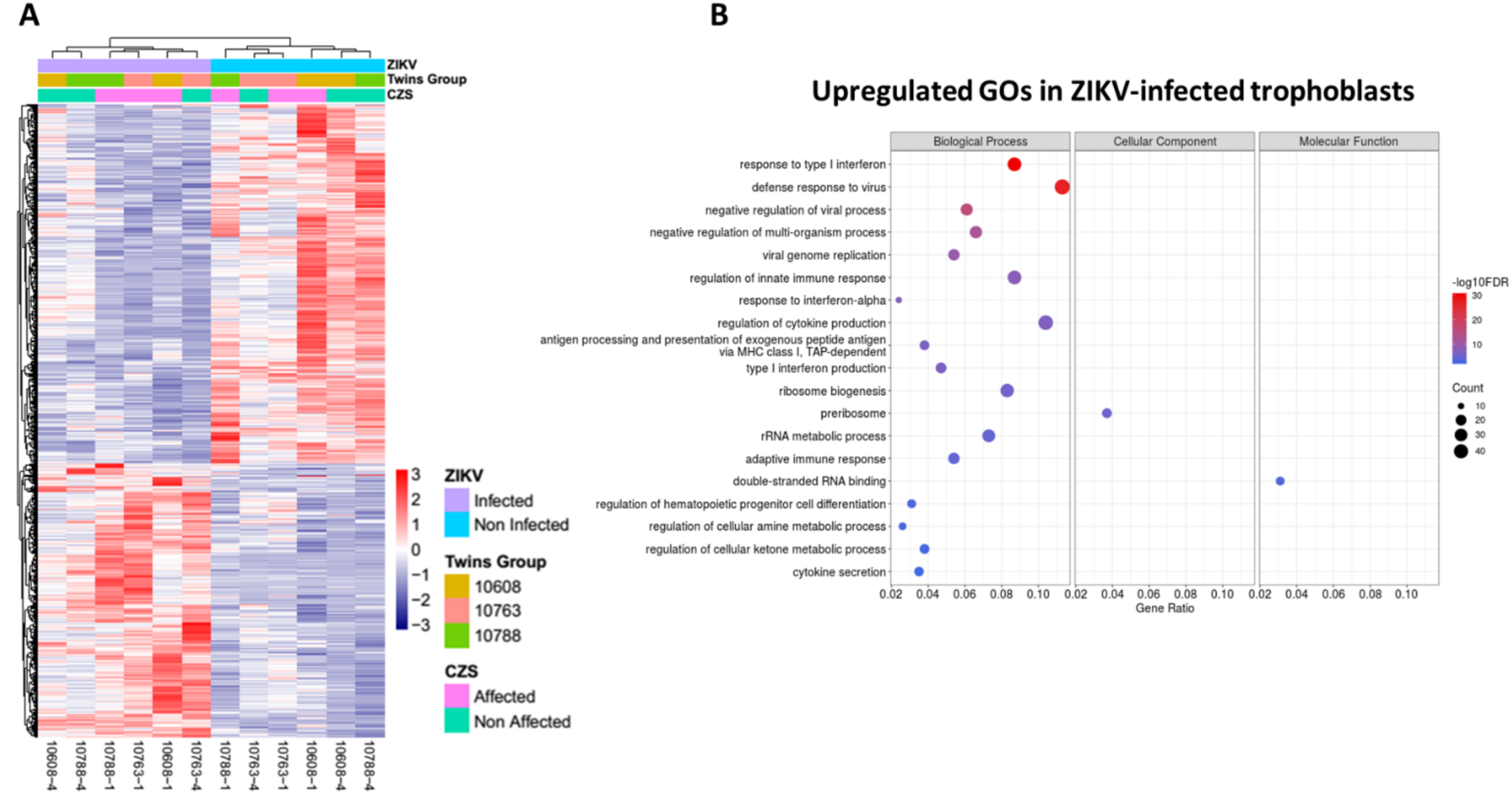
Gene expression analyses of RNA-Seq data from hiPSC-derived trophoblasts from non-affected and CZS-affected twins in culture after ZIKV^BR^ infection. **(A)**. Heatmap representation and clusterization of differentially expressed genes (DEGs) (FDR < 0.05; edgeR exact test), one in each line, in trophoblasts in culture after ZIKV^BR^ infection (purple bar at top) compared with control non-infected trophoblasts (blue bar at top), in cells derived from both non-affected (#10608-4, #10763-4, and #10788-4) and CZS-affected (#10608-1, #10763-1, and #10788-1) twins (one in each column, as indicated at the bottom). Color scale bar at right = Z score. (**B)**. Gene Ontology terms enrichment analysis of upregulated genes in hiPSC-derived trophoblasts after ZIKV^BR^ *in vitro* infection. The three major GO term categories, namely Biological Process, Cellular Component and Molecular Function are separately represented in each panel. The size of the circles is proportional to the number of genes in each significantly enriched category, as indicated at the lower part scales; the colors show the statistical significance of the enrichment, as indicated by the −log10 FDR values that appear in the color-coded scales at the bottom. A GO enrichment significance cutoff of FDR ≤ 0.05 was used.

**Fig 3.**
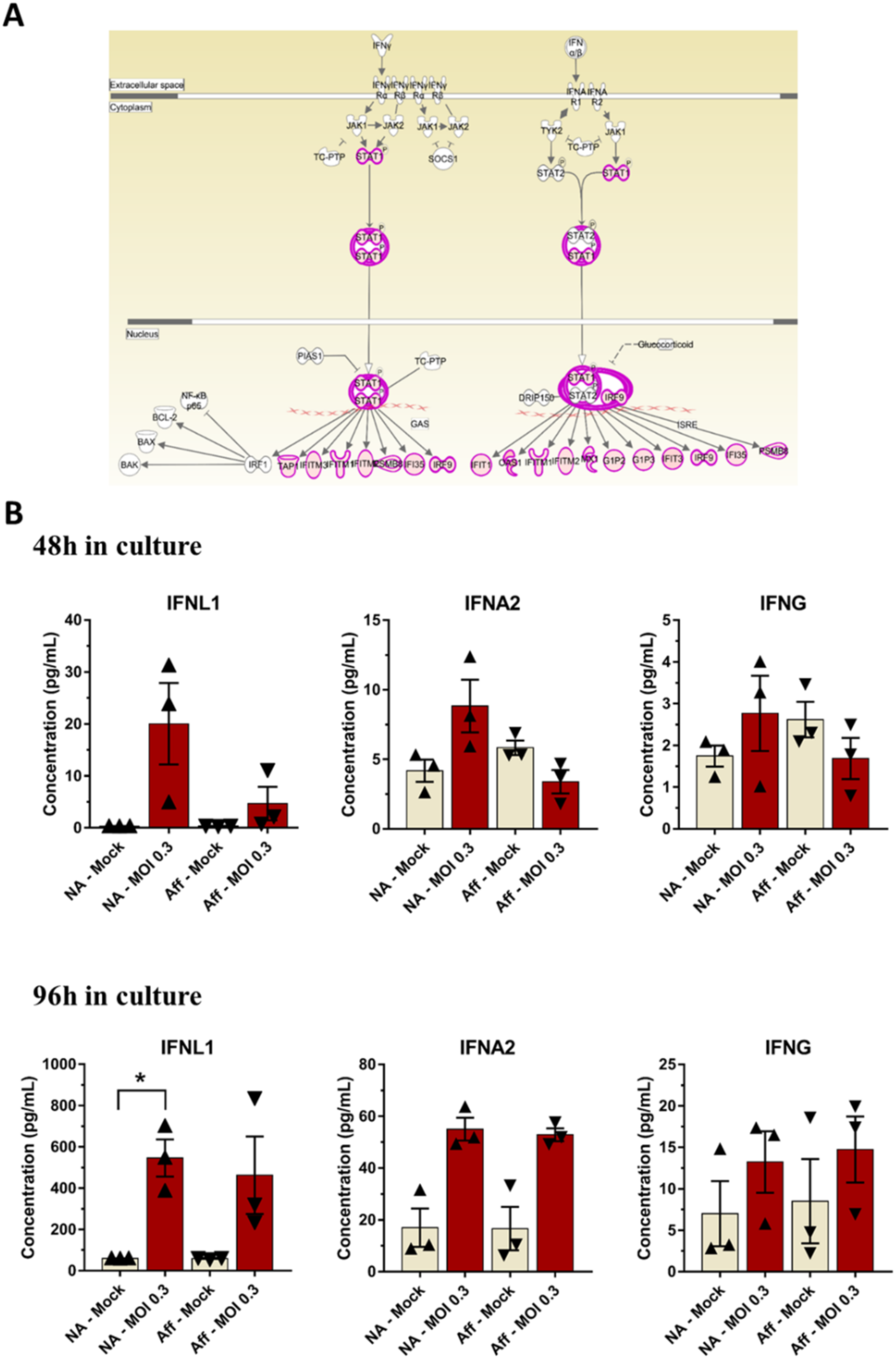
Interferon responses in hiPSC-derived trophoblast after ZIKV^BR^ infection. **(A)**. Pathway enrichment of genes detected as differentially expressed (FDR < 0.05) between hiPSC-derived trophoblasts from non-affected and CZS-affected twins at 96 h after ZIKV^BR^ *in vitro* infection; analysis was carried out with the ingenuity pathway analysis (IPA) tool. Gene upregulation is depicted in shades of red, from white (not significantly changed), to dark red (highly upregulated). **(B)**. Luminex quantitation of IFNA, IFNG and IFNL1 detected in the supernatants of hiPSC-derived trophoblasts in culture, from non-affected or CZS-affected twins at 48 h or 96 h after ZIKV^BR^ infection. Data are represented as mean ± SEM and p-value * < 0.05 (Paired Student’s t test).

We also quantified in the hiPSC-derived trophoblasts the secreted levels of type I (IFNA2), type II (IFNG) and type III (IFNL1) IFNs produced by these cells in the absence of virus or after infection with ZIKV^BR^ and culture for 48 h (**Fig 3B, upper panel**) or 96 h (**Fig 3B, lower panel**). After 48 h in culture without infection, hiPSC-derived trophoblasts from both CZS-affected and non-affected twins were able to secrete IFNA2 and IFNG, whereas IFNL1 was not detectable. There was no statistically significant increase in any IFN secretion at 48 h after ZIKV^BR^ infection of trophoblasts (**Fig 3B, upper panel**). After 96 h in culture, hiPSC-derived trophoblasts from both CZS-affected and non-affected twins secreted higher levels of IFNA2, IFNG and IFNL1 compared with those at 48 h, although high variability in the secreted IFN levels was observed among the trophoblasts from different individuals (**Fig 3B, lower panel**). More importantly, a statistically significant increase in the secretion of IFNL1 by trophoblasts from non-affected twins was observed at 96 h after infection with ZIKV^BR^, but not by trophoblasts from CZS-affected twins (**Fig 3B, lower panel**).

Taken together, these results indicate that hiPSC-derived trophoblasts from both CZS-affected and non-affected twins were able to respond to ZIKV^BR^ infection by secreting IFNA2, IFNG and IFNL1, which induced the upregulation of a set of ISG genes potentially involved in the response to ZIKV^BR^ infection. In addition, only trophoblasts from non-affected twins showed a significant increase in secreted IFNL1 at 96 hpi with ZIKV^BR^ (**Fig 3B, lower panel**).

### A set of genes involved with extracellular matrix organization and leukocyte activation was downregulated in trophoblasts from CZS-affected twins after ZIKV^BR^ infection

We next investigated if there were differences in gene expression between trophoblasts from CZS-affected and non-affected twins after ZIKV^BR^ infection. In total, 44 genes were downregulated after ZIKV^BR^ infection in trophoblasts from CZS-affected when compared with non-affected twins (FDR < 0.05, Edge R exact test) (**Fig 4A and S4 Table**). Different Gene Ontology (GO) terms were found to be significantly (FDR < 0.05, cumulative hypergeometric distribution) enriched among these 44 downregulated DEGs, including “extracellular matrix” and “regulation of leukocyte activation” (**Fig 4B, S5 Table**). Significantly enriched Gene Ontology (GO) terms that were found among the upregulated DEGs are shown in **S6 Fig** and **S5 Table**, and the most significantly enriched categories are “amino acid biosynthetic process” and “acid secretion”.

**Fig 4.**
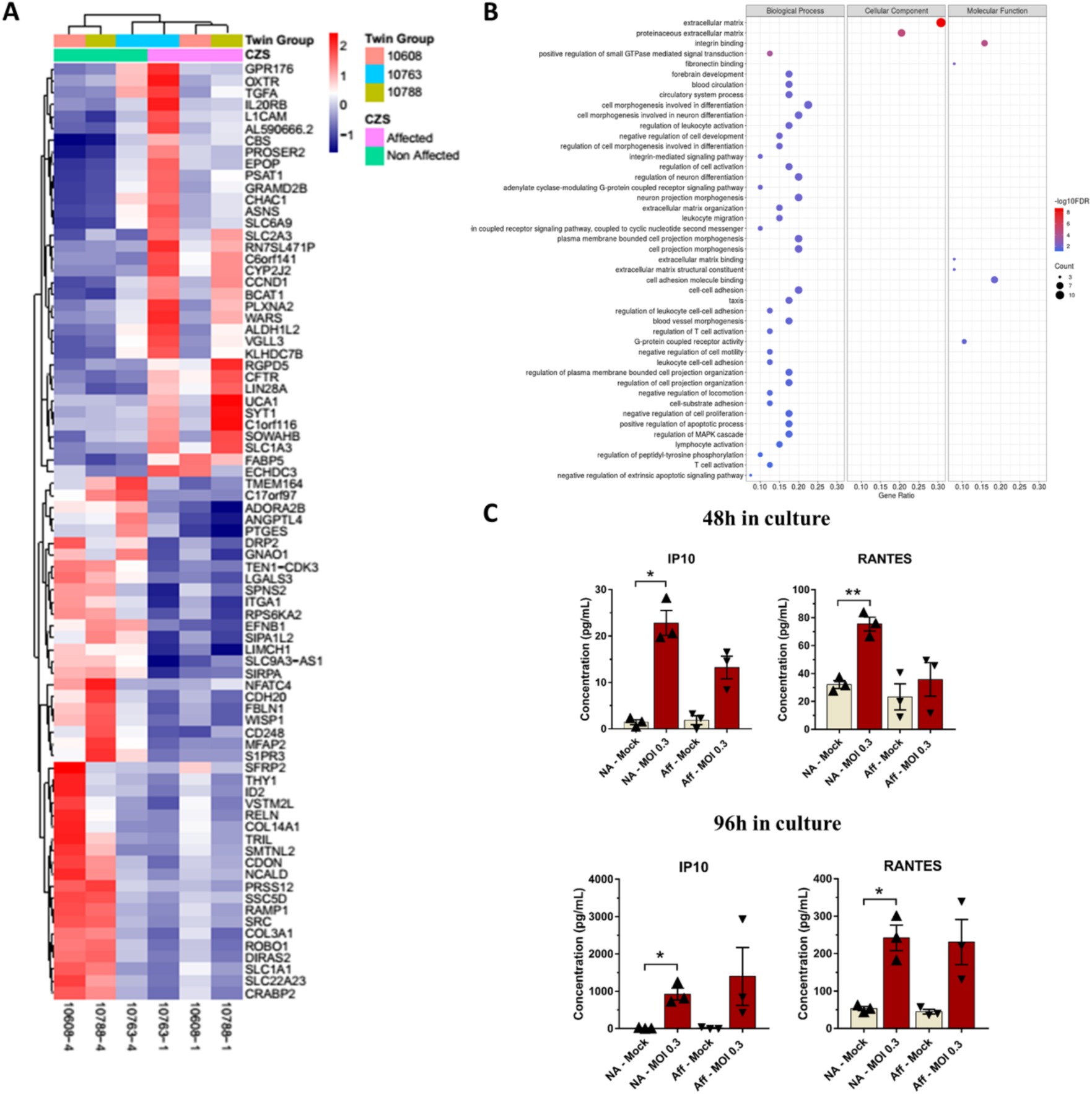
Differential gene expression between hiPSC-derived trophoblast from CZS-affected and non-affected twins after ZIKV^BR^ infection. **(A)**. Heatmap representation and clusterization of differentially expressed genes (DEGs) (FDR < 0.05; edgeR exact test), one in each line, after ZIKV^BR^ *in vitro* infection of trophoblasts from non-affected twins (green bar at top; #10608-4, #10763-4, and #10788-4) as compared with trophoblasts from CZS-affected twins (pink bar at top; #10608-1, #10763-1, and #10788-1) (one in each column, as indicated at the bottom). Color scale bar at right = Z score. (**B)**. Gene Ontology terms enrichment analysis of downregulated genes in hiPSC-derived trophoblasts from CZS-affected twins when compared with trophoblasts from non-affected twins after ZIKV^BR^ *in vitro* infection. The three major GO term categories, namely Biological Process, Cellular Component and Molecular Function are separately represented in each panel. The size of the circles is proportional to the number of genes in each significantly enriched category, as indicated at the scales in the lower part; the colors show the statistical significance of the enrichment, as indicated by the −log10 FDR values that appear in the color-coded scales at the bottom. A GO enrichment significance cutoff of FDR ≤ 0.05 was used. **C**. Luminex quantitation of RANTES/CCL5 and IP10 detected in the culture supernatants of hiPSC-derived trophoblasts from non-affected twins or from CZS-affected twins at 48 h (left panels) or 96 h (right panels) after ZIKV^BR^ infection. Data are represented as mean ± SEM and p-value * < 0.05 (Paired Student’s t test).

The levels of a panel of cytokines and chemokines secreted by the hiPSC-derived trophoblasts were quantified in the supernatants of 48 h or 96 h cell cultures in the absence of virus or after infection with ZIKV^BR^. Interestingly, from all the analytes tested, the chemokines RANTES/CCL5 and IP10 showed a consistent (both at 48 h and 96 h post-infection) and significant increase in secretion by trophoblasts from non-affected twins after infection with ZIKV^BR^, but not by trophoblasts from CZS-affected twins (**Fig 4C**). RANTES/CCL5 secretion levels by trophoblasts from non-affected twins increased 2.4- and 4.6-fold at 48 h or 96 h after infection with ZIKV^BR^, respectively, while IP10 secretion levels increased 16- and 96-fold (**Fig 4C**). The other tested cytokines and chemokines did not show statistically significant differences in the levels produced by infected trophoblasts from CZS-affected or non-affected twins.

## Discussion

Here, we were able to analyze, for the first time, the *in vitro* viral susceptibility and the gene expression patterns after *in vitro* ZIKV^BR^ infection of hiPSC-induced trophoblasts from dizygotic (DZ) twins discordant for the presence of microcephaly. Because these are DZ twins whose mothers were infected with ZIKV^BR^ during pregnancy, the two fetuses in each of the three twin pairs were exposed to the virus at the same time, thus representing a rare and unique cohort to test whether the host genetic background plays any role in determining CZS outcome. Indeed, we have previously shown that hiPSC-induced NPCs from the same subjects exhibit differential *in vitro* ZIKV^BR^ susceptibility, as NPCs from CZS-affected twins had significantly higher ZIKV^BR^ replication and reduced cell growth when compared with NPCs from non-affected twins [32]. No rare Mendelic potentially pathogenic variant was assigned in this cohort that could explain NPCs susceptibility to CZS [32], suggesting that CZS caused by ZIKV may be a multifactorial disorder. Herein, using a well-established model that recapitulates the primitive placenta formed during fetal implantation [34,35], we sought to investigate if hiPSC-induced trophoblasts from the same twins also show differential responses to *in vitro* ZIKV^BR^ infection.

Many studies have established causal effects between ZIKV infection during pregnancy and microcephaly developed in the fetus [15], using *in vitro* as well as mice and non-human primates *in vivo* models [50-54]. Cells localized in the epidermis and dermis were primarily considered as targets for ZIKV infection [55], as mosquito bite remains the major transmission route [56]. ZIKV infection exhibits broad distribution and persistence in body tissues and fluids, and the presence of ZIKV in the amniotic fluid of pregnant women and in semen [57-61] suggests the additional possibility of sexual and perinatal transmissions. Trophectoderm cells of pre-implantation human embryos can be infected with ZIKV [62]. Also, endometrial stromal cells are highly permissive to ZIKV infection, and likely represent a crucial cell target of ZIKV reaching them, either via the uterine vasculature in the viremic phase of the infection or by sexual viral transmission, being a potential source of virus spreading to placental trophoblasts during pregnancy [63]. Indeed, recent studies suggest that the placenta is the key mediator for vertical transmission of ZIKV from infected mothers to fetal brains; several placental cells have been shown to be infected by ZIKV, including placental macrophages, trophoblasts and fibroblasts of the maternal *decidua basalis* [25-29,35,58]. Here, we show that hiPSC-induced trophoblasts from DZ twins discordant for CZS were differentially susceptible to infection with ZIKV^BR^. Interestingly, 96 hpi with ZIKV^BR^, virus titers in culture supernatants were significantly higher in the CZS-affected twins’ trophoblasts when compared with non-affected (**Fig 1C**), indicating that ZIKV^BR^ can replicate more efficiently in the trophoblasts from CZS-affected twins, potentially facilitating virus dissemination into fetal tissues.

Noteworthy, our RNA-seq results indicate that hiPSC-derived trophoblasts express the ZIKV candidate receptor genes (*HSPG2* and *TAM* receptor genes) and the IFN receptor genes, and at lower levels the IFN genes; however none of these genes were differentially expressed between the trophoblasts from CZS-affected and non-affected twins either before or after ZIKV^BR^ infection.

Interestingly, ZIKV^BR^ infection caused a significant increase in IFNL1 secretion by trophoblasts from non-affected twins (**Fig 3B**), whereas in trophoblasts from CZS-affected twins no significant increase was observed (**Fig 3B**). IFNL1 is a type III IFN constitutively released by primary human trophoblasts from full-term placentas [25], which are known to be refractory to ZIKV infection [25,35]. The increased secretion of IFNL1 by trophoblasts from non-affected twins, already at 48 hpi by ZIKV^BR^, but especially at 96 hpi, may protect against ZIKV^BR^ dissemination, as shown in the female reproductive tract of mice [47,64,65]. Our results indicate a possible role of IFNL1 in the control of ZIKV infection by primitive trophoblasts. In addition, after ZIKV^BR^ infection, trophoblasts from the non-affected twins were able to significantly induce the secretion of immune mediator chemokines RANTES/CCL5 [66,67] and IP10 [68], while trophoblasts from the CZS-affected twins were not (**Fig 4C**). RANTES/CCL5 promotes trophoblasts cell migration and can recruit immune cells [69-71], whereas IP10 plays a role as a chemotactic molecule implicated in the migration of trophoblast cells [72]. Thus, lower secretion of RANTES/CCL5 and IP10 indicates that CZS-affected twins’ trophoblasts have a lower ability of migration, immune cell recruitment and viral control.

The placenta plays a critical role in immunological protection and can undergo major structural and functional adaptations in order to protect the fetus from environmental stressors [48,49]. When the placental function is impaired [73], the intrauterine environment might be perturbed and the placental defenses involved in fetal protection compromised. In the RNA-Seq analysis, we found 79 differentially expressed genes in common among the three CZS-affected compared with non-affected twins, 44 of which were downregulated after ZIKV^BR^ infection in the trophoblasts from CZS-affected compared with non-affected twins. The top GO category associated with these downregulated differentially expressed genes is “extracellular matrix”. It is known that for successful fetus development, one of the critical steps is proper invasion of the maternal decidua by trophoblasts [74] and that many molecules, including galectins, are involved in this process [75,76].

One of the downregulated genes in the trophoblasts from CZS-affected twins after ZIKV^BR^ infection was *COL3A1* (**Fig 4A**). Collagens are the main structural proteins in the extracellular matrix, and they have been related to pregnancy and/or placental pathological conditions including gestational diabetes and pre-eclampsia [77,78]. *ITGA1*, another gene downregulated in trophoblasts from CZS-affected twins after ZIKV^BR^ infection, interacts with the extracellular matrix, particularly with collagen and laminin [79]. Importantly, extravillous trophoblasts (EVT) have been shown to express ITGA1 as they invade from the anchoring villi deeply into the maternal endometrium and myometrium in weeks 8–13 of gestation [80,81]. EVTs fail to express ITGA1 in preeclampsia, which is associated with both poor trophoblast invasion and oxidative stress [81,82]. In addition, *LGALS3* gene (encoding galectin-3) was downregulated in trophoblasts from CZS-affected twins after ZIKV^BR^ infection. LGALS3 is a galectin that has been described as involved in the process of trophoblast cell migration and invasion, significant for human embryo implantation [83,84]. Overall, downregulation of genes involved in trophoblast adaptation to the intrauterine environment, including *ITGA1, COL3A1* and *LGALS3* (**S7 Fig**), in the trophoblasts of CZS-affected twins after ZIKV^BR^ infection, may affect trophoblast migration, implantation, homeostasis and possibly impair the control of ZIKV^BR^ infection. Noteworthy, all the above differentially expressed genes were found in common among the three CZS-affected compared with non-affected twins, in spite of the genetic background variability of our cohort. Importantly, as all experiments were done under the same conditions, these differences in genetic background underlie both possible differences in progression of trophoblasts differentiation and the resulting difference in susceptibility to ZIKV infection. Thus, the set of 79 differentially expressed genes found here represents the minimum core of genes significantly altered in common among the three pairs of twins. This study provides genetic targets to be further explored as possible infection susceptibility factors in the placenta.

Our gene expression results indicate that ZIKV^BR^ infection has caused, only in the trophoblasts from CZS-affected twins, the downregulation of genes important for trophoblast adhesion as well as immune response activation. Noteworthy, when gene expression differences between the NPCs from these non-affected and CZS-affected twins were analyzed [32], another set of 64 genes was found to be differentially expressed, including *FOXG1* and *LHX2*, two transcription factors important for neural development [85,86], which were down-regulated in NPCs from CZS-affected compared with non-affected twins [32]. Overall, development of congenital Zika syndrome might result, among other factors, from a concomitant decreased ability of the placenta to respond to ZIKV infection in the CZS-affected neonates, along with a deregulation of neural development genes in ZIKV-infected NPCs of these CZS-affected neonates.

Based on our gene expression analyses, we suggest that the ability to respond more efficiently to ZIKV infection in the placenta may be a key parameter to predict the success of ZIKV dissemination into fetuses’ tissues. Moreover, further understanding of the participation of immune mediators, such as the chemokines RANTES/CCL5 and IP10, in the trophoblast response to ZIKV infection may open a path for drug development or repurposing to possibly inhibit viral replication or avoid viral dissemination into fetus’ tissues.

## Materials and methods

### Human subjects

Three pairs of DZ-D (dichorionic and diamniotic) twins discordant for the presence of microcephaly (#10608-1 and #10608-4; #10763-1 and #10763-4; #10788-1 and #10788-4) whose peripheral blood mononuclear cells had been previously collected and isolated [32] were examined in this study. Zygosity had been previously confirmed by whole-exome sequencing (WES) and by microsatellite analysis [32]. All babies were born from mothers negative for previous STORCH infections and in each affected baby (#10608-1, #10763-1 and #10788-1) the head circumference was three standard deviations (SD) below the mean for the given age, sex, and gestation stage at birth [32,87]. Diagnosis of microcephaly due to ZIKV infection (CZS) in all the three affected subjects was confirmed by neuroimaging, serology and by the mother reporting ZIKV infection symptoms during pregnancy (**S6 Table** and Caires-Junior et al. [32]). The protocol used in this study was approved by the Human Research Ethics Committee from Biosciences Institute, University of São Paulo (protocol # 184/2016). All mothers gave written informed consent in accordance with the Declaration of Helsinki.

### Cell lines and maintenance of hiPSCs

hiPSCs were generated according to Caires-Junior et al. [32]. Briefly, CD71+ cells were isolated from the three pairs of DZ-D twins’ peripheral blood samples (#10608-1 and #10608-4; #10763-1 and #10763-4; #10788-1 and #10788-4). CD71-positive cells were sorted using magnetic labeled antibody (Miltenyi) following the manufacturer’s instructions. All hiPSC lines were tested for ZIKV infection by RT-qPCR using primers described in **S7 Table** and the results were negative. The reprogramming protocol was performed with episomal vectors system (Addgene plasmids 27077, 27078 and 27080), and using the Amaxa human CD34+ cells Nucleofection Kit (Lonza), following the manufacturer’s recommendations. Three days after nucleoporation, cells were seeded on irradiated murine embryonic fibroblasts (Millipore, A24903) in embryonic stem cell (ESC) medium (Dulbecco’s modified Eagle’s medium (DMEM)/F12 supplemented with 2 mM GlutaMAX-I, 0.1 mM non-essential amino acids, 100 μM 2-mercaptoethanol, 20% knockout serum replacement (all provided by Life Technologies), 10 ng/mL bFGF (Peprotech), 0.25 mM NaB, 0.5 mM VPA, 2 μM thiazovivin, 0.5 μM PD 0325901 and 2 μM SB 431542; all provided by Tocris Bioscience). The typical hiPSC colonies were transferred to hESC-qualified Matrigel (Corning)-coated 60 mm petri dishes (Corning) and cultured in Essential 8 Medium (Gibco) with 100 μg/mL normocin (InvivoGen). All derived cell lines were checked for mycoplasma contamination periodically.

### Differentiation of human iPSCs into trophoblasts

Differentiation of human iPSCs into trophoblasts was performed according to Amita et al. [34]. Briefly, hiPSCs were seeded at 15,000 cells/cm^2^ on Matrigel (Corning) coated plates and cultured in mTESR1 medium (Stem Cell Technologies) for 3 days with daily medium changes. Next, basal medium was changed to hES medium supplemented with 4 ng/mL FGF-2 for 24 h. For the next 9 days, hES-BAP medium was used, which consisted of hES medium supplemented with BMP4 (10 ng/mL), A83-01 (1 µM) and PD173074 (0.1 µM), with daily medium changes. hES medium was composed of DMEM/F-12 supplemented with 20 % KO-serum replacement, 1 % NEAA, 1 % glutamine, 0.1 mM 2-mercaptoethanol, all from Gibco.

### In situ immunofluorescence

hiPSCs and trophoblast cultures were fixed with 4 % PFA followed by permeabilization with 0.01 % Triton X-100 and then blocked with 5 % bovine serum albumin (BSA) for 1 h. The cells were incubated overnight with primary antibodies at 4 °C (Anti-hCG beta/β-CG/hCGB3, Abcam #ab53087 and Anti-Cytokeratin 7/CK7/KRT7, Abcam #ab9021), washed with PBS and subsequently incubated with secondary fluorescent antibodies for 1 h at room temperature. Finally, cells were stained with DAPI (4’,6-diamidino-2-phenylindole) and or Phalloidin for 2 min at room temperature. Confocal analysis was performed using Zeiss LSM 800.

### Zika virus

ZIKV^BR^ was a courtesy of Dr. Pedro Vasconcelos [88], Instituto Evandro Chagas, Brazil. Viral stock was established after viral propagation for two serial passages in VERO cells (ATCC® CCL-81™) in serum-free medium (VP SFM, Thermo scientific).

### Infection of the hiPSC-derived trophoblasts

hiPSC-derived day-9 trophoblasts were seeded into 6-well or 24-well plates (Corning) and 2-well chamber slides (Nunc; Thermo Fisher Scientific) to a confluence of 3 × 10^4^ cells/cm^2^. Trophoblasts (monolayer) were exposed to ZIKV^BR^ (MOI: 0.3, 3 and Mock). Monolayer cells were exposed to the virus for 1 h at 37 °C and 5 % CO_2_ (g), washed with hES-BAP medium, and then maintained by up to 96 h (end point).

### Measurement of viral burden

For ZIKV titration, plaque assay was performed with the supernatants of cell cultures. For plaque assay, an amount of 6 × 10^4^ VERO cells/well were seeded in 24-well plates 48 h before the assay. Samples were serially diluted in DMEM culture medium from 10^−1^ to 10^−6^, applied in duplicates of 100 µL to each well, and incubated for 30 min at 37 °C. After virus adsorption, wells were overlaid with culture medium containing carboxymethyl cellulose (1 %) and incubated at 37 °C. After 5 days, plates were drained, washed with PBS, and stained with 0.1 % naphthol blue-black, 1.6 % sodium acetate in 6 % glacial acetic acid for 30 min. Plaque formation units were visually determined in the most appropriate viral dilution and expressed as PFU/mL.

### RNA-Seq assay

Total RNA from hiPSC-derived trophoblasts was extracted using the RNeasy Micro Kit (Qiagen, 74004), treated with TURBO DNase (Ambion, AM2238) for 30 min at 37 °C, and then re-purified with the Qiagen RNeasy Micro Kit. RNA samples were quantified using the Qubit RNA HS Assay Kit (Thermo Fisher Scientific, Q32852); purity was evaluated using NanoDrop ND-1000 Spectrophotometer (NanoDrop Technologies) and the integrity was verified using the Agilent RNA 6000 Pico Kit (Agilent Technologies, 5067-1513) in the 2100 Bioanalyzer Instrument (Agilent Technologies). Stranded tagged cDNA libraries were prepared using the KAPA Stranded mRNA-Seq Kit (Illumina, KK8421) and cluster generation was performed using the Illumina HiSeq 4000 PE Cluster Kit (Illumina, PE-410-1001). Tagged libraries were pooled and sequenced (300 cycles, paired-end sequencing) in the Illumina HiSeq 4000 instrument using a HiSeq 4000 SBS Kit (Illumina, FC-410-1003). Raw reads were preprocessed using the standard Illumina pipeline to segregate multiplexed reads.

### RNA-seq data processing and analysis

Paired-end adapters and low quality reads were removed by fastp version 0.20.0 [89] using the quality-filtering parameters -l 20 −5 3 −3 3. Filtered paired-end reads were mapped and quantified at gene level by the STAR-RSEM pipeline [90,91], using an index made from GRCh38.p12 GencodeV.28 [92]. All differential expression (DE) analyses used the bioconductor package edgeR [93]. To call differentially expressed genes a general linear models (glm) was fitted, and likelihood ratio tests (lrt) were performed using twins’ covariates as blocking groups in all cases. P-values were adjusted using FDR, all genes with FDR lower than 0.05 were considered differentially expressed genes. To identify up-regulated or down-regulated genes, the logCPM data from edgeR were used. Heatmaps were plotted using the R package pheatmap. Enriched gene ontology analysis was perform using the bioconductor package clusterProfiler [94], based on the annotation of the bioconductor package org.Hs.eg.db. All plots were generated with the R packages gg plot2, cowplot and the bioconductor package DOSE.

### Reverse transcription - quantitative PCR (RT-qPCR)

Total RNA from hiPSCs and hiPSC-derived trophoblasts was extracted as described above. The reverse transcription (RT) reaction was performed with 150 ng of each total RNA sample using the SuperScript IV First-Strand Synthesis System (Life Technologies, cat. #18091050) and random hexamer primers in a 20 μL final volume. The obtained cDNAs were diluted 10 times in water and quantitative PCR was performed using 2.5 μL of each diluted cDNA in a total volume of 10 μL containing 1× LightCycler 480 SYBR Green I Master Mix (Roche Diagnostics, cat. #04707516001) and 800 nM of each primer in a LightCycler 480 System (Roche Diagnostics). RT-qPCR was run in two biological replicates with three technical replicates each and primers are shown in **S7 Table**. The GAPDH gene (NM_002046) was used as the reference for internal normalization.

### Cytokine and chemokine protein quantification

Supernatants from trophoblasts’ cell cultures were collected after 48 h or 96 h of infection with ZIKV^BR^ and then evaluated for 33 cytokines/chemokines for the following analytes: IFNalfa2, IFNbeta, IFNg, IFN-lambda1/ IL-29, IFN-lambda 2/ IL28a, IL6, IL10, IL12p40, IL12p70, IL13, IL15, IL17A, IL1RA, IL1a, IL1b, IL2, IL3, IL4, IL5, IL7, IL8, IP10, MCP1, MIP1a, MIP1b, RANTES, TNFa, TNFb, VEGF, EGF, EOTAXIN, GCSF and GMCSF. The Human Cytokine/Chemokine Magnetic Bead Panel kit (Millipore) was used according to the manufacturer’s recommendations. Samples were analyzed in the MagPix instrument (Luminex) and the data was analyzed with the Milliplex Analyst software (Millipore).

### Quantification and statistical analysis

Two-tailed unpaired t test was used for pairwise comparisons. Graphpad Prism software was used to perform statistical analysis (version 7.0). Quantification of data are represented as mean ± SEM and p-value threshold was * 0.05, ** 0.01, *** 0.001 and **** 0.0001.

## Data availability

The RNA-Seq data that support the findings of this study have been deposited in the Sequence Read Archive (SRA) NCBI repository under Accession number PRJNA565997. All other data supporting the findings of this study are available within the article and its Supporting Information files or are available from the authors upon request.

## Acknowledgments

We thank Dr. R. Michael Roberts and Dr. Toshihiko Ezashi, University of Missouri, Columbia, MO, USA for advice on trophoblast cells cultivation protocols. We thank Dr. J.C. Setubal for access to the computational facilities of the Bioinformatics Laboratory of Instituto de Química, Universidade de São Paulo, São Paulo, Brazil.

## Author contributions

Conceptualization: Murilo Sena Amaral, Sergio Verjovski-Almeida

Data Curation: Murilo Sena Amaral, David Abraham Morales-Vicente, Sergio Verjovski-Almeida

Formal Analysis: Murilo Sena Amaral, Ernesto Goulart, Luiz Carlos Caires-Júnior, Alessandra Soares-Schanoski, Jorge E. Kalil, Mayana Zatz, Sergio Verjovski-Almeida

Funding Acquisition: Mayana Zatz, Sergio Verjovski-Almeida

Investigation: Murilo Sena Amaral, Ernesto Goulart, Luiz Carlos Caires-Júnior, Alessandra Soares-Schanoski, Roselane Paiva Gomes, Giovanna Gonçalves de Oliveira Olberg

Methodology: Murilo Sena Amaral, Ernesto Goulart, Luiz Carlos Caires-Júnior, David Abraham Morales-Vicente, Alessandra Soares-Schanoski, Sergio Verjovski-Almeida

Project Administration: Mayana Zatz, Sergio Verjovski-Almeida

Resources: Renato Mancini Astray, Mayana Zatz, Sergio Verjovski-Almeida

Software: David Abraham Morales-Vicente

Supervision: Jorge E. Kalil, Mayana Zatz, Sergio Verjovski-Almeida

Validation: Murilo Sena Amaral, Ernesto Goulart, Luiz Carlos Caires-Júnior, David Abraham Morales-Vicente

Visualization: Murilo Sena Amaral, David Abraham Morales-Vicente

Writing – Original Draft Preparation: Murilo Sena Amaral, Sergio Verjovski-Almeida

Writing – Review & Editing: Murilo Sena Amaral, Ernesto Goulart, Luiz Carlos Caires-Júnior, Alessandra Soares-Schanoski, Renato Mancini Astray, Jorge E. Kalil, Mayana Zatz, Sergio Verjovski-Almeida

## Supporting information

**S1 Fig.**
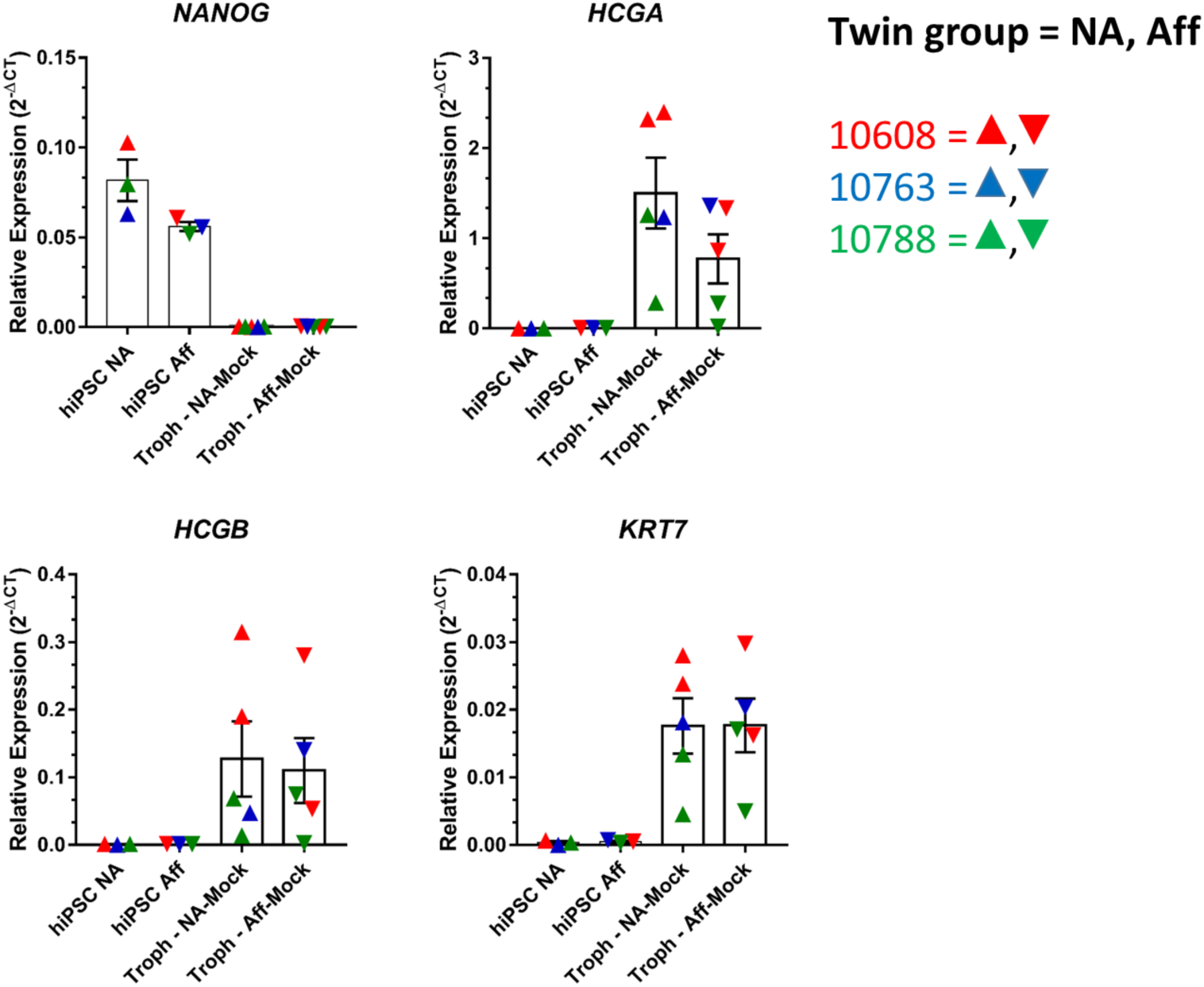
Expression measured by RT-qPCR of marker genes of hiPSCs and trophoblasts. Expression measured by RT-qPCR of *NANOG*, a hiPSC marker, and of *HCGA, HCGB* and *KRT7*, three of the genes upregulated in the trophoblasts as compared with the hiPSCs. Twins from each family are represented with a different color: red, #10608 twins; blue, #10763 twins; green, #10788 twins. (mean ± SEM; n = 2 biological replicates, except for #10763 due to sample loss during culture)

**S2 Fig.**
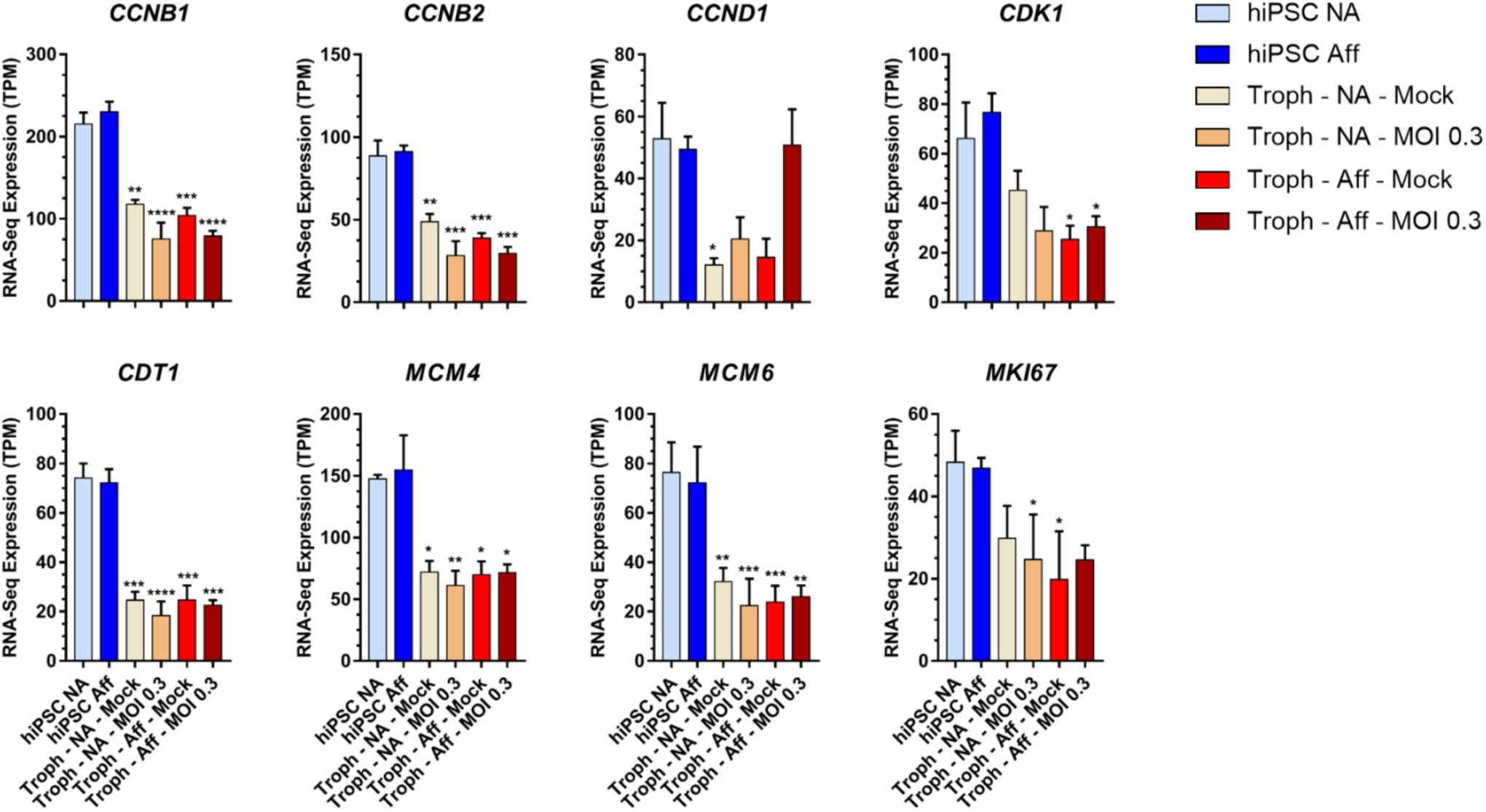
Expression levels measured by RNA-Seq of proliferation-related genes in the hiPSCs and in the hiPSC-derived trophoblasts from non-affected or CZS-affected twins. Related to Figure 2. The bars represent expression levels (in TPM) of selected genes associated with cellular proliferation in hiPSCs from non-affected (light blue, hiPSC NA) or CZS-affected (dark blue, hiPSC Aff) twins, in the hiPSC-derived trophoblasts from non-affected twins’ mock (yellow, Troph – NA-Mock) or ZIKV-infected cells (orange, Troph – NA-MOI 0.3), and in the hiPSC-derived trophoblasts from CZS-affected twins’ mock (red, Troph – Aff-Mock) or ZIKV-infected cells (brown, Troph – Aff-MOI 0.3). The levels of expression of the genes were compared between non-affected (both non-infected and infected) trophoblasts and non-affected hiPSCs; and from CZS-affected (both non-infected and infected) trophoblasts and CZS-affected hiPSCs. Genes significantly down-regulated in trophoblast cells when compared with hiPSCs are shown (one-away ANOVA, p-value threshold was * 0.05, ** 0.01, *** 0.001 and **** 0.0001). Error bars show SEM.

**S3 Fig.**
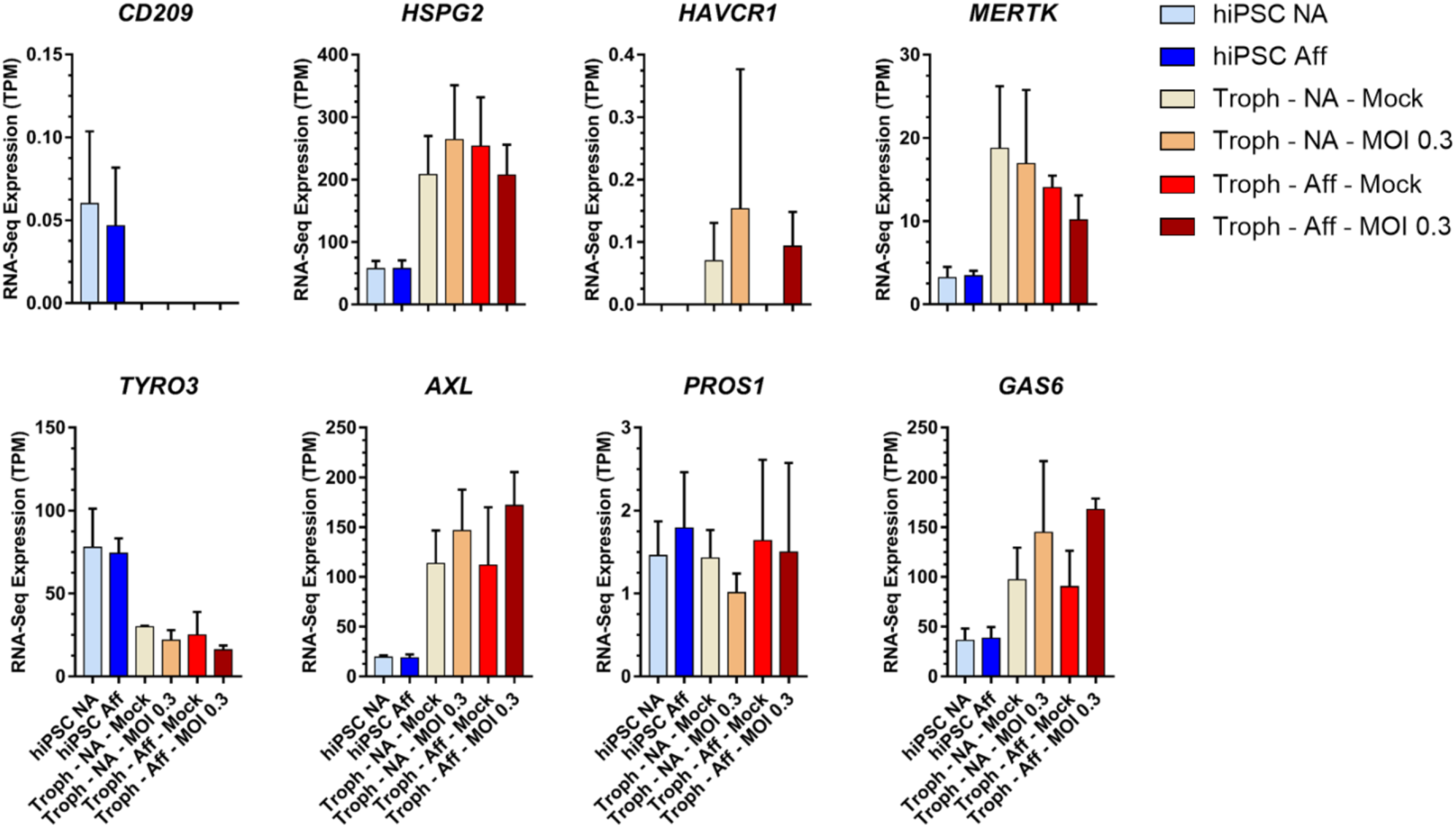
Expression levels measured by RNA-Seq of genes encoding candidate attachment factors implicated in ZIKV infection of human cells. Related to Figure 2. The bars represent expression levels (in TPM) of eight candidate ZIKV attachment factor genes in hiPSCs from non-affected (light blue, hiPSC NA) or CZS-affected (dark blue, hiPSC Aff) twins, in the hiPSC-derived trophoblasts from non-affected twins’ mock (yellow, Troph – NA-Mock) or ZIKV-infected cells (orange, Troph – NA-MOI 0.3), and in the hiPSC-derived trophoblasts from CZS-affected twins’ mock (red, Troph – Aff-Mock) or ZIKV-infected cells (brown, Troph – Aff-MOI 0.3). None of these genes was significantly differentially expressed in hiPSC-derived trophoblasts from CZS-affected twins, when compared with hiPSC-derived trophoblasts from non-affected twins in pairwise comparisons (two-tailed t-test, equal variance). Error bars show SEM.

**S4 Fig.**
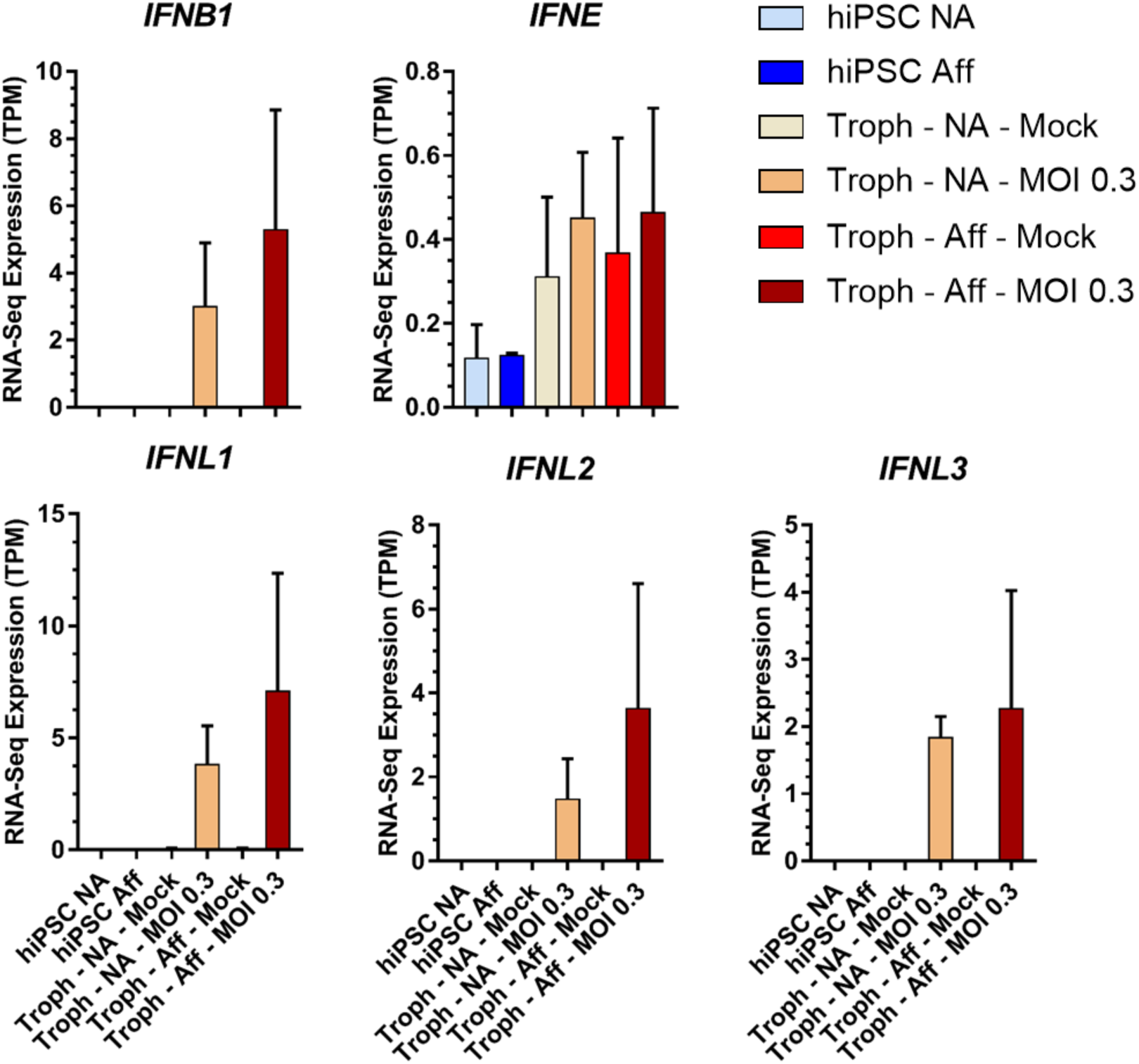
Expression levels measured by RNA-Seq of genes encoding representatives of Type I, Type II, and Type III IFNs. Related to Figure 2. The bars represent expression levels (in TPM) of genes encoding interferons in hiPSCs from non-affected (light blue, hiPSC NA) or CZS-affected (dark blue, hiPSC Aff) twins, in the hiPSC-derived trophoblasts from non-affected twins’ mock (yellow, Troph – NA-Mock) or ZIKV-infected cells (orange, Troph – NA-MOI 0.3), and in the hiPSC-derived trophoblasts from CZS-affected twins’ mock (red, Troph – Aff-Mock) or ZIVK-infected cells (brown, Troph – Aff-MOI 0.3). None of these genes was significantly differentially expressed in hiPSC-derived trophoblasts from CZS-affected twins when compared with hiPSC-derived trophoblasts from non-affected twins in pairwise comparisons (two-tailed t-test, equal variance). None of the 13 recognized human *IFNA* (*IFNA1, 2, 4, 5, 6, 7, 8, 10, 13, 14, 16, 17, 21*), *IFNG, IFNK* and *IFNW1* genes were significantly expressed in any of the data sets. Error bars show SEM.

**S5 Fig.**
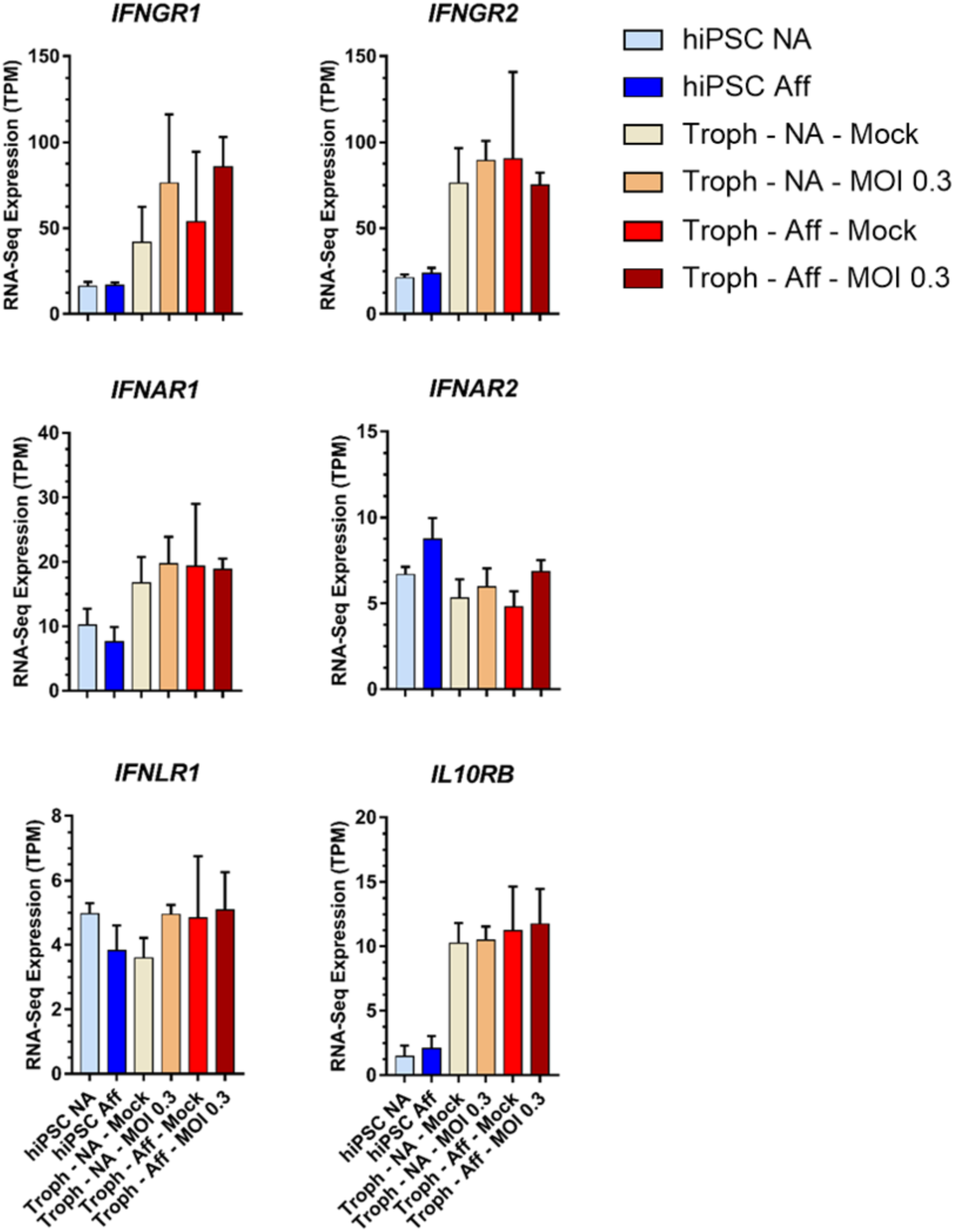
Expression levels measured by RNA-Seq of genes encoding receptors for Type I, Type II, and Type III IFNs. Related to Figure 2. The bars represent expression levels (in TPM) of genes encoding interferon receptors in hiPSCs from non-affected (light blue, hiPSC NA) or CZS-affected (dark blue, hiPSC Aff) twins, in the hiPSC-derived trophoblasts from non-affected twins’ mock (yellow, Troph – NA-Mock) or ZIKV-infected cells (orange, Troph – NA-MOI 0.3), and in the hiPSC-derived trophoblasts from CZS-affected twins’ mock (red, Troph – Aff-Mock) or ZIKV-infected cells (brown, Troph – Aff-MOI 0.3). None of these genes was significantly differentially expressed in hiPSC-derived trophoblasts from CZS-affected twins when compared with hiPSC-derived trophoblasts from non-affected twins in pairwise comparisons (two-tailed t-test, equal variance). Error bars show SEM.

**S6 Fig.**
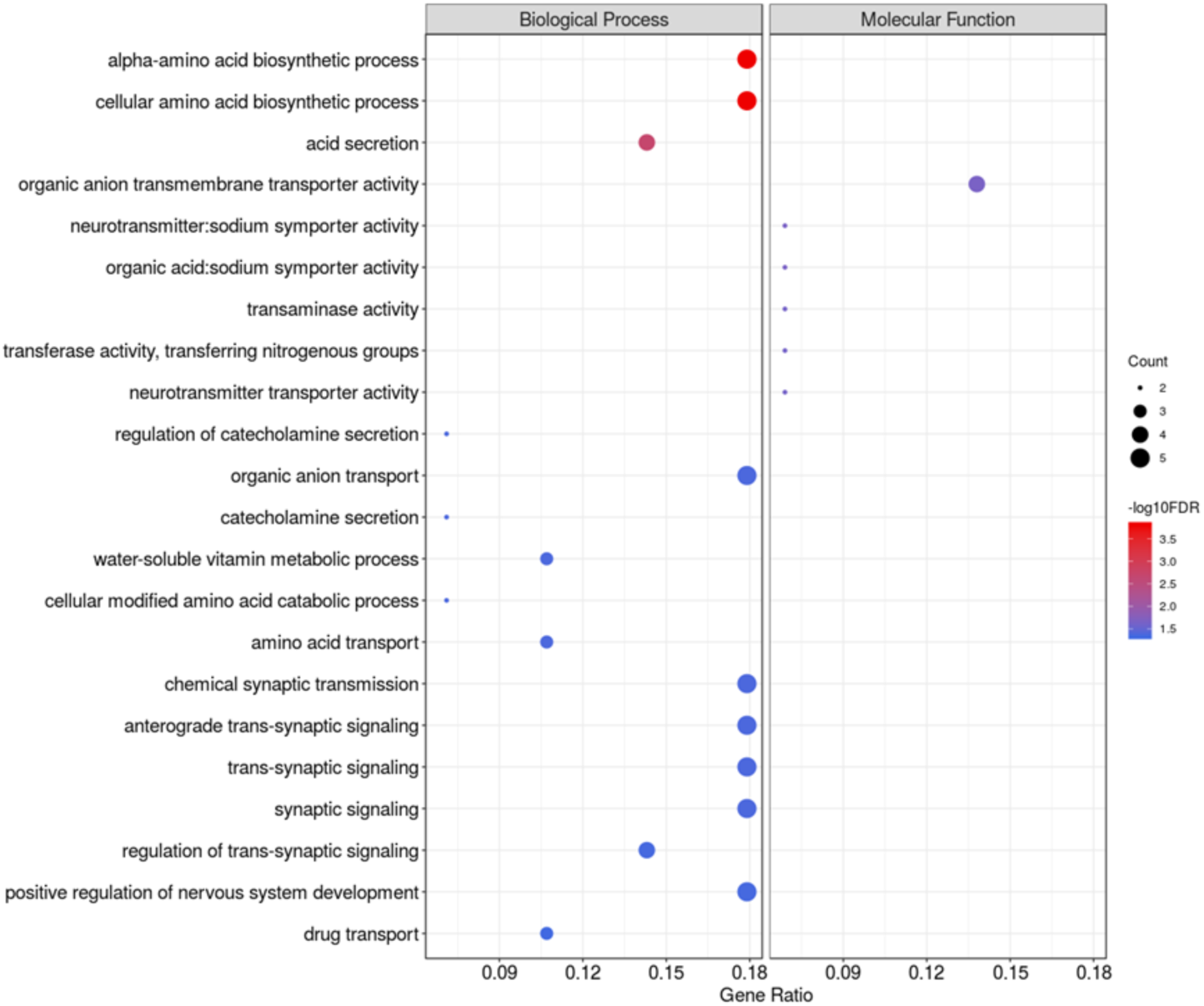
Differential gene expression between hiPSC-derived trophoblast from CZS-affected and non-affected twins after ZIKV^BR^ infection. Related to Figure 4. Gene Ontology terms enrichment analysis of upregulated genes in hiPSC-derived trophoblasts from CZS-affected compared with non-affected twins after ZIKV^BR^ *in vitro* infection. The major GO term categories, namely Biological Process and Molecular Function are separately represented in each panel. The size of the circles is proportional to the number of genes in each significantly enriched category, as indicated at the lower part scales; the colors show the statistical significance of the enrichment, as indicated by the −log10 FDR values that appear in the color-coded scales at the bottom. A GO enrichment significance cutoff of FDR ≤ 0.05 was used.

**S7 Fig.**
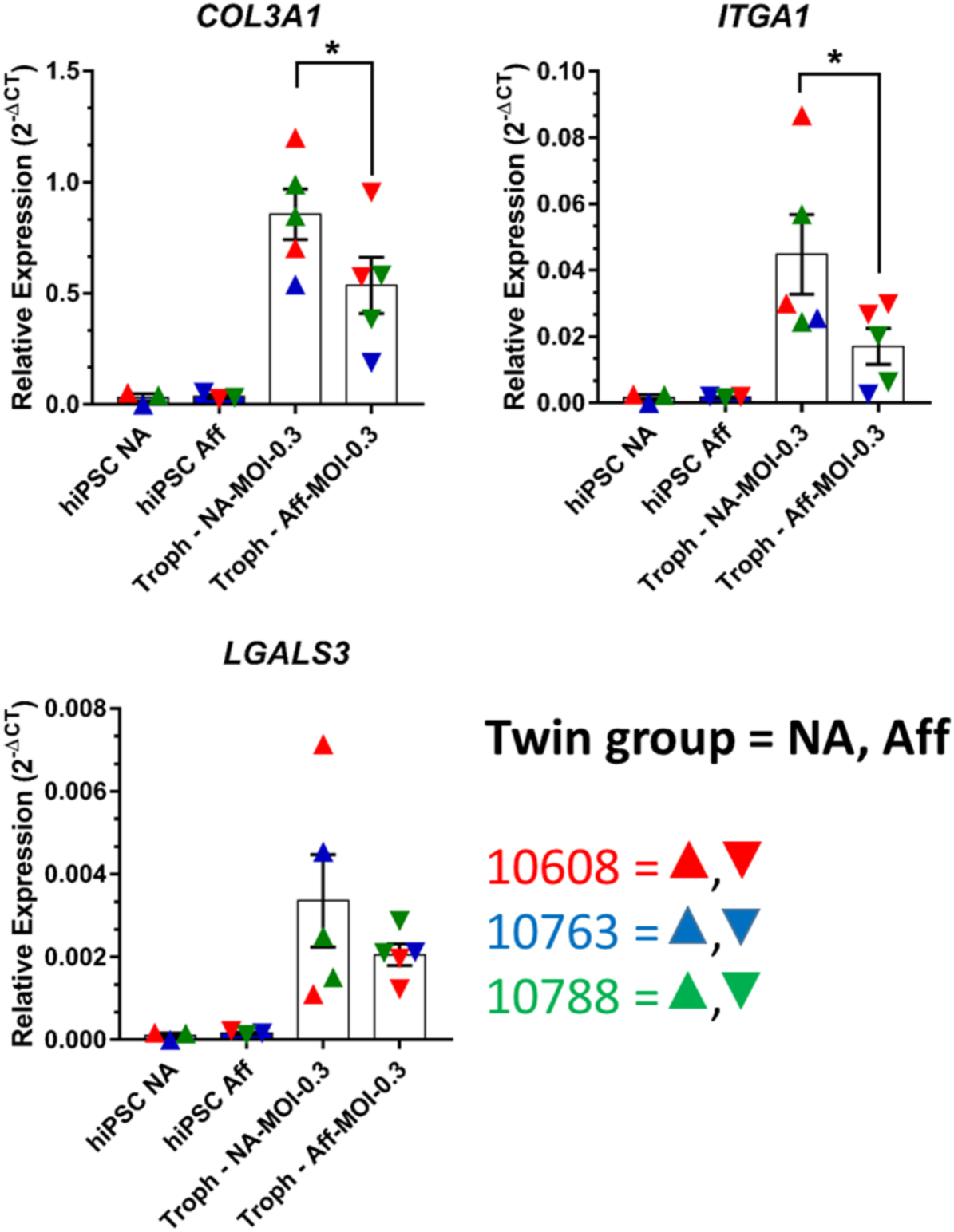
Expression measured by RT-qPCR of genes found in the RNA-Seq analysis downregulated after ZIKV^BR^ infection in trophoblasts from CZS-affected when compared with non-affected twins. Expression measured by RT-qPCR of *COL3A1, ITGA1* and *LGALS3*, three of the genes downregulated after ZIKV^BR^ infection in trophoblasts from CZS-affected (Aff) when compared with non-affected (NA) twins. Twins from each family are represented with a different color: red, #10608 twins; blue, #10763 twins; green, #10788 twins. Mean ± SEM is shown. (n = 2 biological replicates, except for #10763 due to sample loss during culture; One-tailed t-test, * p<0.05).

